# Regulation of global transcription in *E. coli* by Rsd and 6S RNA

**DOI:** 10.1101/058339

**Authors:** Avantika Lal, Sandeep Krishna, Aswin Sai Narain Seshasayee

## Abstract

In *Escherichia coli*, the sigma factor σ^70^ directs RNA polymerase to transcribe growth-related genes, while σ^38^ directs transcription of stress response genes during stationary phase. Two molecules hypothesized to regulate RNA polymerase are the protein Rsd, which binds to σ^70^, and the non-coding 6S RNA which binds to the RNA polymerase- σ^70^ holoenzyme. Despite multiple studies, the functions of Rsd and 6S RNA remain controversial. Here we use RNA-Seq in five phases of growth to elucidate their function on a genome-wide scale. We show for the first time that Rsd and 6S RNA facilitate σ^38^ activity throughout bacterial growth, while 6S RNA also regulates widely different genes depending upon growth phase. We discover novel interactions between 6S RNA and Rsd and show widespread expression changes in a strain lacking both regulators. Finally, we present a mathematical model of transcription which highlights the crosstalk between Rsd and 6S RNA as a crucial factor in controlling sigma factor competition and global gene expression.

## INTRODUCTION

In bacteria, the core RNA polymerase (α_2_ββ′ω, referred to as E) transcribes all cellular genes. Proteins called sigma (σ) factors bind to E forming Eσ holoenzymes, and direct it to transcribe from specific promoters. *Escherichia coli* has seven sigma factors (Ishihama 2000) – σ^70^ (RpoD), σ^38^ (RpoS), σ^32^, σ^54^, σ^28^, σ^24^ and σ^19^. Though it was previously reported that the cellular concentration of E exceeded that of sigma factors (Jishage and Ishihama 1995; Maeda *et al.* 2000), recent quantitation has shown that sigma factors exceed E under common culture conditions (Grigorova *et al.* 2006; Piper *et al.* 2009), and may compete to bind to E. This is supported both by *in vitro* assays demonstrating competition between sigma factors under limiting E (Maeda *et al.* 2000; Jishage *et al.* 2002; Laurie *et al.* 2003; Bernardo *et al.* 2006) and *in vivo* studies where increasing activity of one sigma factor decreased the activity of others and vice-versa (Farewell *et al.* 1998; Jishage *et al.* 2002; Laurie *et al.* 2003).

σ^70^ (RpoD) is the housekeeping sigma factor, directing transcription of genes essential for growth. The major alternative sigma factor, σ^38^ (RpoS), is present in low concentration during exponential growth, but regulates several hundred genes (Rahman *et al*. 2006; Dong *et al*. 2008). Upon entry into stationary phase, while the concentrations of E and σ^70^ show little change, σ^38^ accumulates (Gentry *et al.* 1993; Tanaka *et al.* 1993; Piper *et al.* 2009; Table 1) and directs transcription of genes for multiple stress tolerance (Lange and Hengge-Aronis 1991; Hegge-Aronis *et al.* 1991; Hengge-Aronis and Fischer 1992, Weber *et al.* 2005). However, σ^70^ remains the most abundant sigma factor, and has the highest affinity for E (Maeda *et al*. 2000; Colland *et al.* 2002; Piper *et al.* 2009; Ganguly and Chatterji 2012), implying that additional regulators are needed for σ^38^ to compete effectively with σ^70^.

**Table 1:** Cellular concentrations of RNA polymerase, sigma factors, Rsd and 6S RNA

| Molecule | Exponential | Stationary | Reference |
| --- | --- | --- | --- |
| Core RNA Polymerase | ~4.3 $\mu\text{M}$ | ~4.3 $\mu\text{M}$ | Piper *et al.* 2009 |
| $\sigma^{70}$ | ~12.1 $\mu\text{M}$ | ~12.0 $\mu\text{M}$ | |
| $\sigma^{38}$ | Not Detected | ~2.7 $\mu\text{M}$ | |
| Rsd | ~5.5 $\mu\text{M}$ | ~10.4 $\mu\text{M}$ | |
| 6S RNA | ~1000 molecules/cell | ~10,000 molecules/cell | Wassarman and Storz 2000 |

An example of such a regulator is the Crl protein, which binds to σ^38^ and increases its affinity for E, and promotes transcription by the Eσ^38^ holoenzyme at some promoters (Bougdour et al. 2004; Typas et al. 2007; England et al. 2008). The nucleotide ppGpp also increases the ability of alternative sigma factors to compete with σ^70^ (Jishage et al. 2002). However, ppGpp is produced transiently on entry into stationary phase (Cavanagh et al. 2010), and Crl has been shown to decrease in extended stationary phase (Bougdour et al. 2004).

On the other hand, two regulators - the protein Rsd and the non-coding 6S RNA - act on σ^70^. Rsd binds to σ^70^, sequestering it from E, and inhibits E σ^70^-dependent transcription at several promoters in vitro (Jishage and Ishihama 1998). An Rsd null strain showed increased transcription from a σ^70^ dependent promoter, and reduced transcription from a σ^38^ dependent promoter, in stationary phase, while Rsd overexpression had the opposite effect (Jishage and Ishihama 1999). It was hypothesized that in stationary phase, Rsd reduces Eσ^70^ formation, and, by freeing E to bind σ^38^, increases Eσ^38^ formation. However, a microarray experiment found no significant change in gene expression on knocking out Rsd, and even comparison of the knockout with an Rsd overexpressing strain found changed expression of only a few σ^38^-dependent genes (Mitchell *et al.* 2007). Also, effects of Rsd were observed only during stationary phase, supposedly due to low Rsd levels in exponential phase. However, Rsd was later observed to be present at ~50% of the level of σ^70^ in exponential phase and ~90% in stationary phase (Piper *et al.* 2009) - raising the question of why no change in expression was seen in its knockout.

6S RNA is a non-coding RNA expressed from the *ssrS* gene, which binds to the Eσ^70^ holoenzyme (Wassarman and Storz 2000). It has been shown to block Eσ^70^ binding to a target promoter (Wassarman and Saecker 2006) and inhibit transcription from several promoters in vitro and in vivo (Wassarman and Storz 2000; Trotochaud and Wassarman2004; Gildehaus et al. 2007; Cavanagh et al. 2008; Cavanagh et al. 2010). A 6S RNA knockout showed increased expression from some Eσ^70^ promoters containing extended-10 elements, and reduced expression from a few Eσ^38^ promoters, in stationary phase(Trotochaud and Wassarman 2004). It was suggested that sequestration of Eσ^70^ by 6S RNA allows σ^38^ to compete more effectively for E, increasing transcription by Eσ^38^. An alternative hypothesis was that 6S RNA regulates a trans-acting factor important for Eσ^38^ activity. There have been two microarray-based genome-wide studies on 6S RNA knockouts. While the first (Cavanagh et al. 2008) showed that 6S RNA regulated hundreds of genes in stationary phase, and an extended-10 element and weak-35 element could make a promoter sensitive to 6S RNA, the second (Neusser et al. 2010) found no correlation of 6S RNA sensitivity with promoter sequence or sigma factor preference. There was also little overlap between the regulated genes found in the two experiments.

Rsd and 6S RNA are present at high concentrations during growth, increase in stationary phase, and are thereafter at high levels (Jishage and Ishihama 1998; Wassarman and Storz 2000; Piper et al. 2009). Yet their regulatory function remains controversial and studies have been contradictory. Both are hypothesized to reduce Eσ^70^ and increase Eσ^38^ formation in stationary phase. However, it is not clear whether Rsd affects the expression of σ^70^ target genes, nor whether it has any effect during growth - and if not, why not? Similarly, with 6S RNA, while studies concur that it reduces transcription from several σ^70^ promoters, one study found that it regulated σ^38^ target genes during stationary phase whereas another did not. Does it have such an effect? If so, is it a direct effect of Eσ^70^ sequestration or mediated by a trans-acting factor? What is that trans-acting factor? Finally, although both Rsd and 6S RNA have been proposed to help σ^38^ compete with σ^70^, one sequesters σ^70^ while the other sequesters the Eσ^70^ holoenzyme. What impact does this have on their regulatory effects? Why does the cell require two molecules to sequester σ^70^?

Here, we present a genome-wide investigation of the functions of Rsd and 6S RNA in *E. coli.* We used RNA-seq to identify genes regulated by Rsd and 6S RNA in five phases of growth and demonstrated that both regulate global transcription during exponential as well as stationary phase. We showed that both increase expression of σ^38^ target genes, with 6S RNA also regulating hundreds of σ^70^ targets, varying greatly with growth phase. We found evidence of substantial crosstalk between Rsd and 6S RNA, with each regulating the other’s expression and non-additive effects on over a thousand genes. Finally, we developed a mathematical model of sigma factor competition in *E. coli*, which explains our experimental results and highlights crosstalk between Rsd and 6S RNA as a vital factor controlling sigma factor competition.

## MATERIALS AND METHODS

**Growth conditions:** Luria-Bertani broth and agar (20 g/L) were used for routine growth. M9 defined medium (0.6% Na_2_HPO_4_, 0.3% KH_2_PO_4_, 0.05% NaCl, 0.01% NH_4_Cl, 0.1 mM CaCl_2_, 1 mM MgSO_4_, 5 × 10^−4^% Thiamin) supplemented with 0.5% glucose and 0.1% casamino acids was used for RNA-seq and validation. During strain construction, ampicillin or kanamycin were used at final concentrations of 100 μg/ml and 50 μg/ml respectively.

**Strain construction:** Single gene deletions were achieved by λ Red recombination (Datsenko and Wanner 2000), using plasmids pKD46 and pKD13 and specific primers. This introduced a kanamycin resistance cassette into the chromosome. Knockout strains were selected on LB Kanamycin plates. In the *rsd* knockout, the resistance cassette was removed by FLP-mediated site-specific recombination using plasmid pCP20. The Δ*rsd*Δ*ssrS* double knockout was generated by P1 transduction from single knockouts (Thomason *et al.* 2007). The 3x-FLAG epitope was added to the C-terminus of Rsd by PCR using plasmid psub11 as template (Uzzau *et al.* 2001) and introduced onto the MG1655 chromosome by λ Red recombination using specific primers. The *ssrS* knockout was moved into this strain using P1 transduction. Strain constructions were verified by PCR using specific primers and Sanger sequencing. Primer sequences are given in Table S4.

**RNA extraction and mRNA enrichment:** Overnight cultures in M9 glucose were inoculated in 100 mL fresh M9 glucose to a final OD_600_ of 0.02 and incubated at 37 °C with shaking. Two biological replicates were performed for each strain. Cells were collected by centrifugation at the early exponential (OD_600_ ~0.3), mid-exponential (OD_600_ ~0.8), transition to stationary (OD_600_ ~1.6), stationary (16 hrs, OD_600_ ~2), and late stationary (48 hrs, OD_600_ ~1.6) phases of growth. RNA was extracted using TRIzol (Invitrogen), following the manufacturer’s protocol. Total RNA was treated with DNase I (Invitrogen, 18068-015) according to the manufacturer’s protocol. Further precipitation of RNA and ribosomal RNA cleanup was achieved using the MICROBExpress bacterial mRNA purification Kit (Ambion, AM1905) according to the manufacturer’s protocol. RNA concentration was determined using a Nanodrop 2000 (Thermo Scientific) and quality was checked by visualization on agarose gels.

**RNA-Seq:** Sequencing libraries were prepared using TruSeq RNA sample preparation kit v2 (Illumina, RS-122-2001) according to the manufacturer’s guidelines, checked for quality on an Agilent 2100 Bioanalyzer, and sequenced for 50 cycles from one end on an Illumina HiSeq1000 platform at the Centre for Cellular and Molecular Platforms, Bangalore. RNA-Seq data is summarized in Table S2.

**qRT-PCR for RNA-Seq validation:** qRT-PCR was carried out using specific primers to selected mRNA targets (primers in Table S4, results in Table S3). 5 ng RNA was used for each RT-PCR reaction. TAKARA One-step SYBR PrimeScript RT-PCR kit II (RR086A) was used according to the manufacturer’s protocol, on an Applied Biosystems ViiA 7 Real-Time PCR system.

**Western Blotting:** Cells were grown as for RNA-seq. For stationary phase samples, 5 ml culture was harvested by centrifugation. For mid-exponential phase, 10 ml was harvested. Lysates were prepared and protein concentration was estimated using BCA assay (Thermo Fisher Scientific, 23227). Lysates containing equal amounts of protein were loaded onto an SDS-PAGE gel. Proteins were elecroblotted onto a nitrocellulose membrane and probed with mouse primary antibody against the protein of interest followed by horseradish peroxidase-conjugated anti-mouse secondary antibody. The primary antibodies were: Mouse monoclonal antibody to RpoB (Neoclone, WP023), Mouse monoclonal antibody to σ^70^ (Neoclone, WP004), Mouse monoclonal antibody to σ^38^ (Neoclone, WP009), Mouse monoclonal anti-FLAG antibody (Sigma-Aldrich, F3165), Mouse monoclonal antibody to GroEL (Abcam, ab82592). Bands were visualized using SuperSignal West Dura Chemiluminescent substrate (Thermo Fisher Scientific, 34076) and imaged using an ImageQuant LAS 4000 system (GE Healthcare Life Sciences). Band intensities were quantified using ImageJ (http://imagej.nih.gov/ij).

**Data sources:** The *E. coli* K-12 MG1655 genome was downloaded from NCBI (NC_000913.2). Gene coordinates were taken from RegulonDB v8.0 (Salgado *et al.* 2013). Lists of σ^38^ target genes and ppGpp regulated genes were obtained from (Weber *et al.* 2005) and (Traxler *et al.* 2008) respectively. Lists of genes regulated by 6S RNA were obtained from (Cavanagh *et al.* 2008) and (Neusser *et al.* 2010). Coordinates of RNA polymerase binding regions and their occupancy were obtained from (Cho et al.2014) A list of 501 genes under constitutive σ^70^ promoters was obtained from (Shimada *et al.* 2014). Of these, we selected 270 genes which were not regulated by σ^38^ or other sigma factors according to RegulonDB (Salgado *et al.* 2013), (Weber *et al.* 2005), or our RNA-Seq from the σ^38^ knockout.

**Data Analysis:** RNA-Seq reads were mapped to the *E. coli* K-12 MG1655 genome using BWA^52^, and reads mapping uniquely were used for further analysis. DESeq (Anders and Huber 2010) was used for differential expression analysis. Fold changes and p-values for all genes are given in the Supplementary Dataset. Genes with <= 10 reads mapping to them under all conditions were excluded from all analyses and plots. All statistical analyses were performed in R version 3.0.1.

**Mathematical model:** The reactions in Figure 6A are represented by a set of differential equations that determine how the dynamical variables (levels of sigma factors, holoenzymes, etc.) change with time. These equations are given in File S2. Since the timescales on which these reactions occur is much faster than typical timescales of cell division, or processes such as stress responses in stationary phase, we assume that the levels of sigma factors, holoenzymes, etc., are in quasi steady-state in the cell. Parameters that determine the specific steady-state include the dissociation constants of the various complexes, total levels of Rsd, 6S RNA, E, sigma factors, and some others, listed in Table 2 with their default values. These values are altered in specific simulations as described in the results.

**Table 2:** Default values of parameters used for simulations. Wherever possible, values are specific to stationary phase. These values are discussed in File S1.

| Parameter | Meaning | Value | Reference |
| --- | --- | --- | --- |
| Volume | Average cell volume | $10^{-15}$ L | (Ali Azam *et al.* 1999) |
| $E_{\text{total}}$ | Total cellular concentration of RNA Polymerase | 4.3 $\mu\text{M}$ | (Piper *et al.* 2009) |
| $\sigma^{70}_{\text{total}}$ | Total cellular concentration of $\sigma^{70}$ | 12.0 $\mu\text{M}$ | |
| $\sigma^{38}_{\text{total}}$ | Total cellular concentration of $\sigma^{38}$ | 2.7 $\mu\text{M}$ | |
| $Rsd_{total}$ | Total cellular concentration of Rsd | 10.4 μM | |
| $6S\ RNA_{total}$ | Total cellular concentration of 6S RNA | 13 μM | (Wassarman and Storz 2000) |
| $DNA_{total}$ | Nonspecific binding sites per cell | $4.6 \times 10^6$ | MG1655 genome size |
| $P_{70}, P_{38}$ | σ-cognate promoters per cell | 200 | (Mauri and Klumpp 2014) |
| $K_{E\sigma 70}$ | Dissociation constant for E - $\sigma^{70}$ binding | 3.3 nM | (Colland *et al.* 2002) |
| $K_{E\sigma 38}$ | Dissociation constant for E - $\sigma^{38}$ binding | 15.2 nM | |
| $K_{NS}$ | Dissociation constant for nonspecific binding of RNA Polymerase and DNA | $10^{-4}$ M | (deHaseth *et al.* 1978) |
| $K_{Rsd}$ | Dissociation constant for Rsd - $\sigma^{70}$ binding | 32 nM | (Sharma and Chatterji 2008) |
| $K_{6S}$ | Dissociation constant for 6S RNA - $E\sigma^{70}$ binding | 131 nM | (Beckmann *et al.* 2011) |
| $K_{E\sigma P}$ | Dissociation constant for holoenzyme - promoter binding | $10^{-7}$ M | (deHaseth *et al.* 1998); (Bintu *et al.* 2005) |
| Operon length | Average operon length | 1000 nt | RegulonDB (Salgado *et al.* 2013) |
| $c$ | Rate of promoter clearance | $0.005\ s^{-1}$ | (Smith and Sauer 1996); (Mauri and Klumpp 2014) |
| $E$ | Rate of escape from elongation | $0.021\ s^{-1}$ | (Proshkin *et al.* 2010) |

The model of holoenzyme formation (shaded in Figure 6A) describes the following reactions:

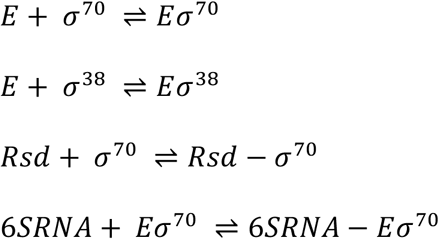

In steady state, the following equations must be fulfilled:

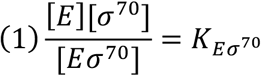

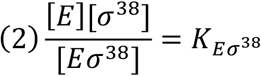

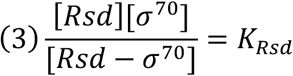

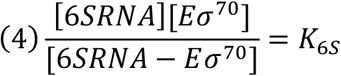

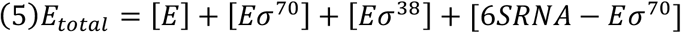

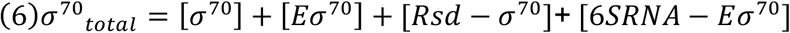

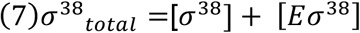

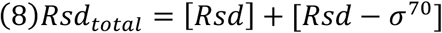

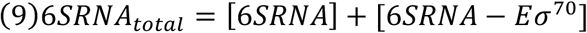

Steady-state levels of the dynamical variables were obtained from the above equations both by solving them numerically and by integrating the corresponding differential equations until they reached steady-state.

The model with DNA (represented by the full schematic in Figure 6A) includes, in addition, the following reactions:

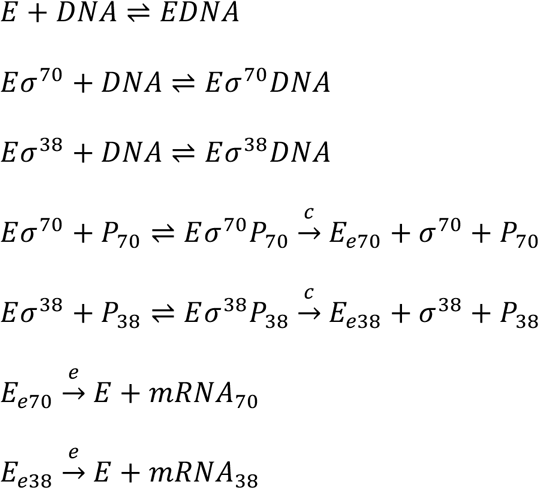

*c* represents the rate of promoter clearance and *e* represents the rate of transcript elongation. At steady state these fulfil the following equations, in addition to equations (1) - (4) and (8) - (9). K_NS_ represents the dissociation constant for non-specific binding of RNA polymerase to DNA, which we assume is equal for E, Eσ^70^ and Eσ^38^. We also assume the specific binding constants of Eσ^70^ and Eσ^38^ to their target promoters to be equal in stationary phase.

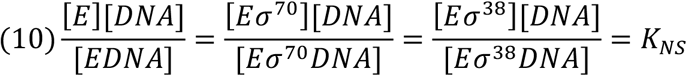

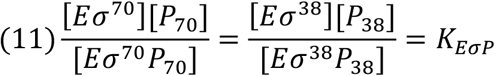

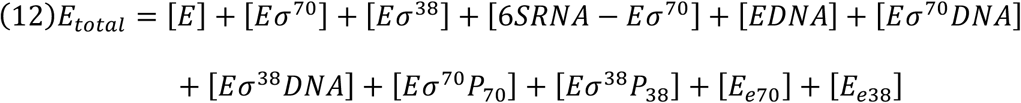

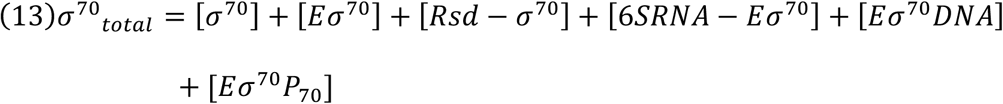

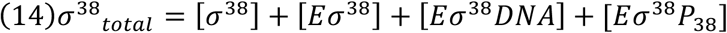

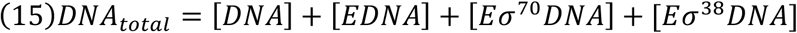

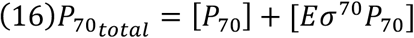

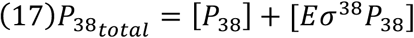

Here, steady-state levels were obtained by integrating the corresponding differential equations until they reached steady-state.

**Data availability:** RNA-Seq data are deposited with NCBI GEO under the accession number GSE74809. Supplementary material has been uploaded to Figshare. These include Figures S1-S12, Tables S1-S4, and Files S1-S3.

## RESULTS

**RNA-seq to identify genes regulated by Rsd and 6S RNA:** Figure 1A shows a schematic of the binding activity of 6S RNA and Rsd. Rsd sequesters σ^70^ and prevents it from binding to core RNA polymerase (E), while 6S RNA binds to the Eσ^70^ holoenzyme and prevents it from binding promoters. To identify their effects on gene expression, we carried out RNA-seq to identify genes regulated by Rsd and / or 6S RNA in five growth phases.

**Figure 1.**
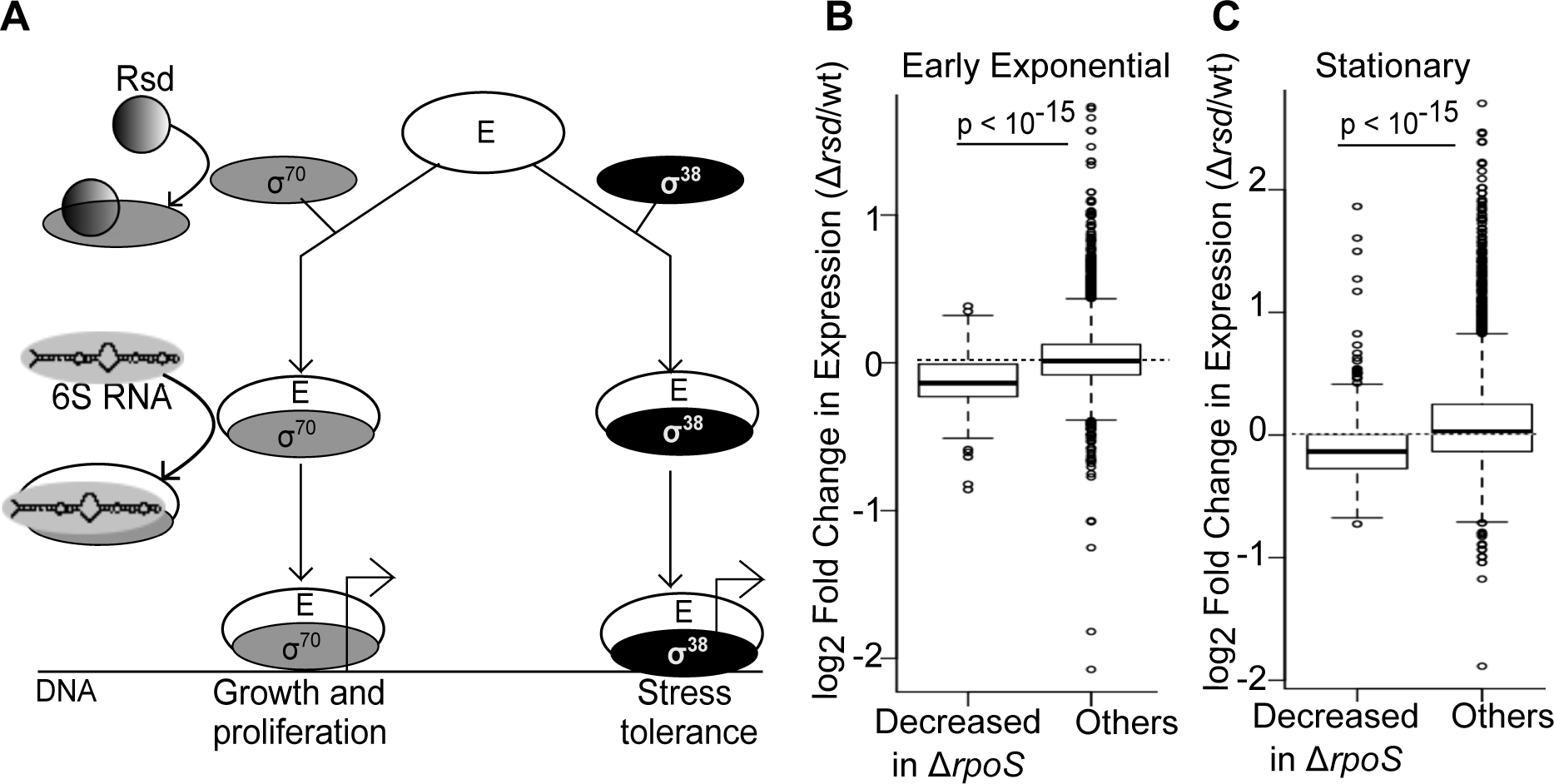
Rsd regulates RpoS target genes throughout growth. (A) Schematic showing the binding activity of Rsd and 6S RNA. (B) Boxplot of log2 fold change in gene expression (Δ*rsd*/wild-type, EE phase) for 313 genes whose expression is reduced at least twofold in Δ*rpoS*/wild-type in EE phase, compared to all other genes. (C) Boxplot of log2 fold change in gene expression (Δ*rsd*/wild-type, stationary phase) for 634 genes whose expression is reduced at least twofold in Δ*rpoS*/wild-type in stationary phase, compared to all other genes, p-values are for Wilcoxon Test.

Table S1 lists strains and plasmids used. Five strains: *E. coli* K-12 MG1655 (Wild-type), Rsd knockout (Δ*rsd*), 6S RNA knockout (Δ*ssrS*), 6S RNA-Rsd double knockout (Δ*rsd*Δ*ssrS)*, and σ^38^/RpoS knockout (Δ*rpoS)* were used for RNA-seq. These had similar growth rates in M9 glucose (Figure S1). RNA-seq was performed in five growth phases: early exponential (EE), mid-exponential (ME), transition to stationary (TS), stationary (S), and late stationary (LS; time points in Methods).

**Rsd increases σ^38^/RpoS activity throughout growth:** We defined differentially expressed genes as genes whose expression changed >=2-fold in a mutant strain relative to wild-type, with FDR-adjusted p-value < 0.05. Using these criteria, the Δ*rsd* strain showed only 16 differentially expressed genes. These included several non-coding RNAs *(ryfD, sokA, oxyS, sroH, sibD)* which were increased two- to six-fold in Δ*rsd* during stationary phase. Notably, 6S RNA expression was altered in Δ*rsd;* 6S RNA was 2.3 times its wild-type level in stationary phase, but in ME phase was reduced to about half its wild-type level.

As very few genes were differentially expressed in Δ*rsd*, we looked for smaller changes. We found that in all growth phases, genes whose expression was significantly reduced (>=2-fold, p < 0.05) in Δ*rpoS* also displayed reduced expression in Δ*rsd.* The box plots in Figure 1B-C show the distribution of log_2_ fold change in gene expression in Δ*rsd* relative to wild-type. ~75% of genes whose expression was significantly reduced in Δ*rpoS* showed reduced expression (log_2_ fold change < 0) in Δ*rsd.* Similar plots for other growth phases are in Figure S2A. This reduced expression was less than twofold in magnitude and so was not seen when searching for differentially expressed genes.

Though these fold changes are small (the median decrease in expression is 10%), we consider them important for several reasons. First, they represent a consistent and highly significant (Wilcoxon test p < 10^−15^ in all growth phases except LS) decrease in expression across hundreds of *rpoS* regulated genes, in all five growth phases. Second, the average overlap between the set of *rpoS* regulated genes in successive growth phases is only 54% - so it is not a single set of genes whose expression is reduced in Δ*rsd*, but a different set in each growth phase. Third, the Δ*rsd* samples showed high inter-replicate correlation in all growth phases (Table S2). Fourth, this trend of decreased expression held true when only previously reported σ^38^ targets (Weber *et al.* 2005) were considered (Figure S2B). Fifth, genes whose expression was increased >=2-fold in Δ*rpoS* also showed increased expression in Δ*rsd*, during EE, ME and stationary phases (Figure S2C). On the other hand, expression of genes controlled by constitutive σ^70^ target promoters (Shimada *et al.* 2014) was not substantially altered in any phase (Figure S3).

Thus, the Rsd knockout behaved like a σ^38^ knockout, only with smaller changes in gene expression. Western blots in stationary phase showed that σ^38^ protein levels were similar in Δ*rsd* and wild-type (Figure S4; σ^38^ level in Δ*rsd* is 94% of wild-type); however, it is extremely difficult to obtain reliable measures of small differences in protein level using western blots. We cannot be certain based on this experiment whether the reduced expression of σ^38^ targets in Δ*rsd* is due to reduced σ^38^ protein, reduced E-σ^38^ association, or both. We attempt to address this question theoretically later.

**6S RNA regulates distinct sets of genes in all phases of growth:** What role does the Eσ^70^ - sequestering 6S RNA play in gene regulation? Is its function similar to that of the σ^70^-sequestering Rsd? Our RNA-seq showed that the 6S RNA knockout (Δ*ssrS*) was very different from the Rsd knockout. To begin with, it showed >=2-fold differential expression of 447 genes in total.

Figure. 2A and B show that there was very little overlap between genes differentially expressed in Δ*ssrS* in successive growth phases, even in the two stationary phase time points. Only a few genes were differentially expressed in Δ*ssrS* throughout growth, particularly *fau*(*ygfA*), which is downstream of 6S RNA in the same operon. *fau* expression was increased when 6S RNA was overexpressed in a wild-type background (Figure S5), indicating that this is at least partly a regulatory effect of 6S RNA and not merely a polar effect.

**Figure 2.**
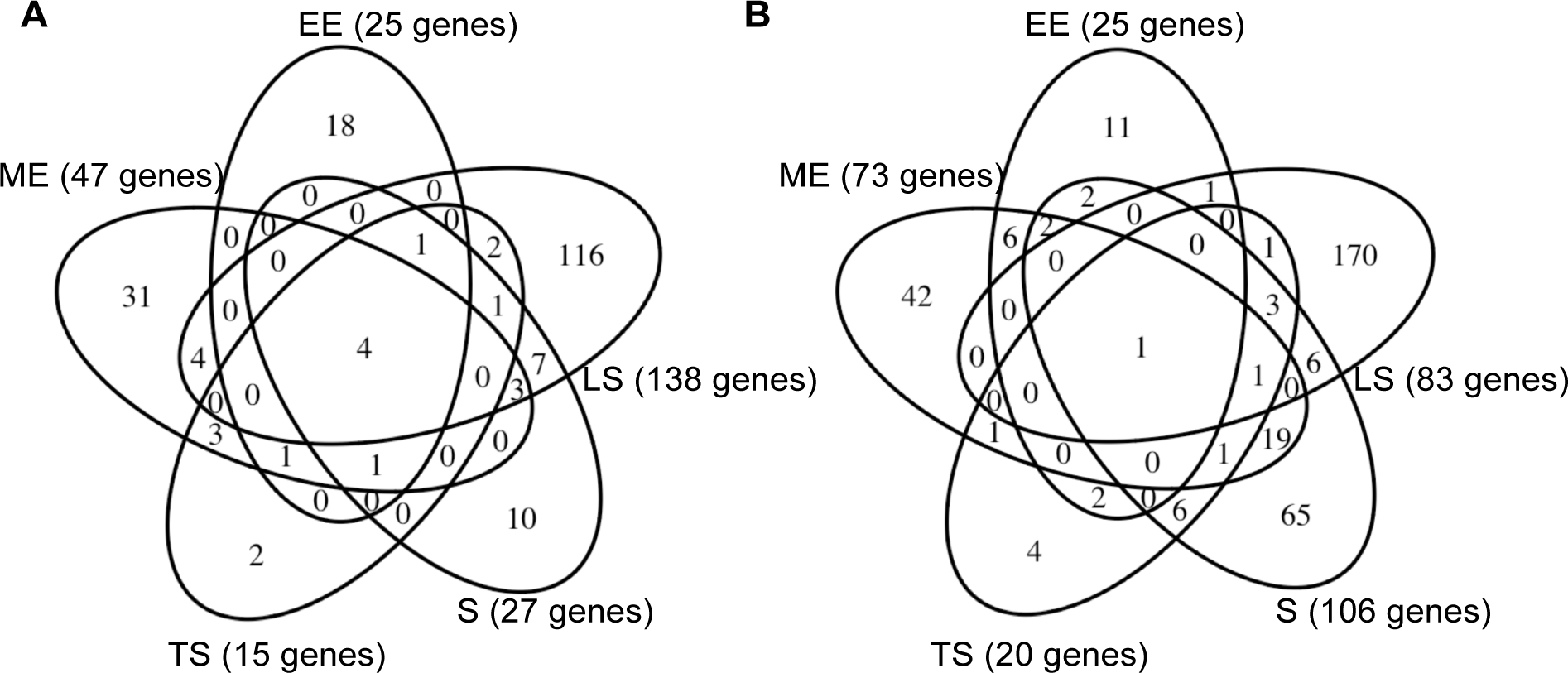
The 6S RNA regulon is highly growth-phase dependent. (A) Venn diagram showing genes whose expression is increased at least twofold in *ΔssrS*, in different phases of growth. (B) Venn diagram showing genes whose expression is reduced at least twofold in *ΔssrS*, in different phases of growth.

During exponential phase, expression of several genes for amino acid transporters (*artM, artI, hisP, hisQ, hisJ, hisM, tdcC*) and amino acid biosynthesis(*argH, argB, thrA, thrB, thrC, asnB, glyA*) was increased in Δ*ssrS*, while expression of genes involved in stress responses (*rmf, appY, yadC, ybcM, yciF, gadW, ydeI, yodD, dps, hdeA, hslV, oppA, osmE, dosC*) was reduced. On the other hand, in LS phase, genes involved in global transcriptional regulation(*crp, hha*) and the TCA cycle (*sdhD, sdhC, gltA, aceB, ppc, sucA*) were upregulated. Thus, the physiological function of 6S RNA appears largely growth phase-dependent; nevertheless, there are certain patterns throughout growth.

**6S RNA increases σ^38^/RpoS activity throughout growth:** Like *Δrsd*, the *ΔssrS* strain showed reduced expression of σ^38^ target genes (Figure. 3A and B, Figure S6, Wilcoxon test p < 10^−15^ in all growth phases except LS). This was not due to reduced σ^38^ protein, as western blots in stationary phase showed that the *ΔssrS* strain had higher σ^38^ protein than the wild-type (Figure S4). On the other hand, expression of genes under constitutive σ^70^ promoters was slightly increased in the ME, TS and LS phases (Figure S7).

**Figure 3.**
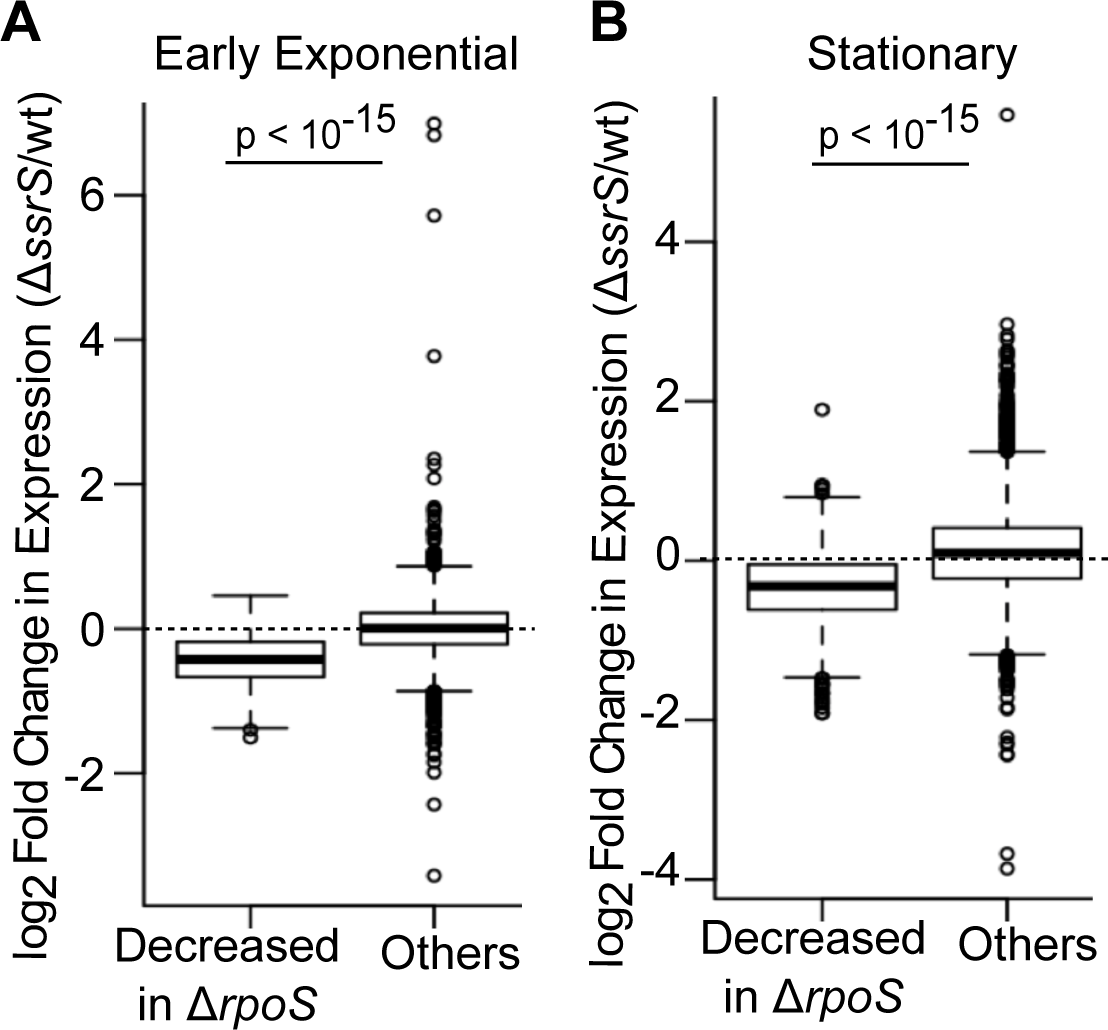
6S RNA regulates RpoS target genes throughout growth. (A) Boxplot of log_2_ fold change in gene expression (Δ*ssrS*/wild-type, EE phase) for 313 genes whose expression is reduced at least twofold in Δ*rpoS*/wild-type in EE phase, compared to all other genes. (B) Boxplot of log2 fold change in gene expression (Δ*ssrS*/wild-type, stationary phase) for 634 genes whose expression is reduced at least twofold in Δ*rpoS*/wild-type in stationary phase, compared to all other genes. p-values are for Wilcoxon Test.

**6S RNA regulates RpoB, Crl and Rsd:** Since the wild-type expression of 6S RNA increased in successive growth phases (Figure 4A), its effect should increase with growth phase. Indeed, with the exception of the TS phase, the number of 6S RNA regulated genes increased with growth phase. A previous study (Cavanagh *et al.* 2010) showed increased ppGpp in a 6S RNA knockout during the TS phase, and we observed consistent changes in gene expression (Figure S8). Since ppGpp also favors competition of alternative sigma factors with σ^70^ (Jishage *et al.* 2002), increased ppGpp may reduce the effects of *ΔssrS* deletion during this phase.

**Figure 4.**
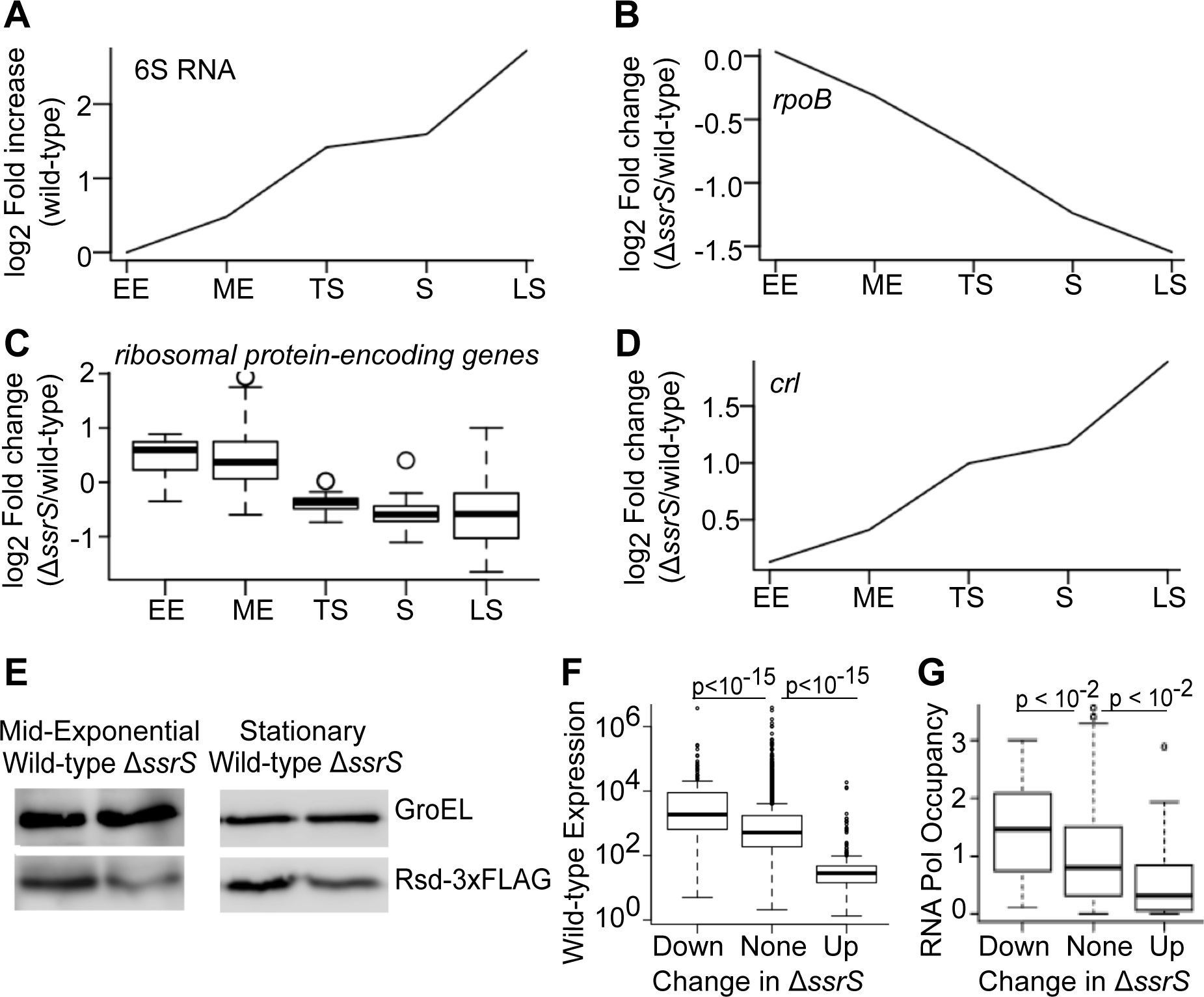
6S RNA controls RpoB, Crl and Rsd. (A) log2 fold increase in 6S RNA levels in wild-type *E. coli* over successive growth phases (relative to its expression in EE phase), based on RNA-Seq. (B) log2 fold change in *rpoB* expression in Δ*ssrS*/wild-type, in successive growth phases. (C) log2 fold change in the expression of 46 ribosomal protein genes in Δ*ssrS*/wild-type, in successive growth phases. (D) log2 fold change in *crl* expression in Δ*ssrS*/wild-type, in successive growth phases. (E) Western blot for 3xFLAG-tagged Rsd in wild-type and Δ*ssrS* backgrounds, in mid-exponential and stationary phases. (F)

We found 36 genes where 6S RNA had a dose-dependent effect throughout growth, increasing in each successive growth phase (Figure S9). For instance, expression of *rpoB* (encoding the β subunit of core RNA polymerase) was reduced in Δ*ssrS*, and the magnitude of reduction increased with growth phase (Figure 4B). Reduced RpoB was validated in stationary phase by qRT-PCR (Table S3) and western blotting (Figure S10). Since excess Eσ^70^ inhibits *rpoB* transcription (Fukuda *et al.* 1978), we suggest that deleting 6S RNA leads to higher free Eσ^70^, which proportionally represses *rpoB.* Since RpoB is limiting for the formation of core RNA polymerase (Piper *et al.* 2009), this implies that the cell compensates for 6S RNA loss by reducing RNA polymerase synthesis. Similarly, we observed decreased expression of genes encoding ribosomal proteins (Figure 4C), consistent with previous reports in stationary phase (Neusser *et al.* 2010).

Boxplots showing the stationary phase wild-type expression level of genes that are reduced >=2-fold in Δ*ssrS*/wild-type, genes that are not differentially expressed in Δ*ssrS*, and genes that are increased >=2- fold in Δ*ssrS*/wild-type. (G) Boxplots showing the stationary phase RNA polymerase occupancy (by ChlP-chip) of promoters belonging to genes that are reduced >=2-fold in Δ*ssrS*/wild-type, genes that are not differentially expressed in Δ*ssrS*, and genes that are increased >=2-fold in Δ*ssrS*/wild-type. Only genes that were first in their transcription unit and were associated with a single RNA polymerase binding site, were included in (G). All p-values are for Wilcoxon Test.

6S RNA also represses some genes in a dose-dependent manner. Figure 4D shows that *crl* expression is increased in Δ*ssrS*, and the magnitude of this effect increases with growth phase. As with the effect of reducing RNA polymerase synthesis to compensate for the loss of 6S RNA, increasing σ^38^ protein and Crl may be a means to compensate for reduced Eσ^38^ activity in Δ*ssrS* bacteria.

Since we observed that Rsd regulated 6S RNA expression, we checked whether 6S RNA in turn regulated Rsd. Our RNA-Seq showed that *rsd* was not differentially expressed in Δ*ssrS.* To check if 6S RNA regulated Rsd post-transcriptionally, we added a 3xFLAG tag to the C-terminal of the Rsd protein, as in previous studies (Piper *et al.* 2009). Indeed, western blots showed that 3xFLAG-tagged Rsd was reduced in the Δ*ssrS* background, in both ME and stationary phases (Figure 4E).

Finally, though 6S RNA binds to Eσ^70^, its effects are highly promoter-specific (Trotochaud and Wassarman 2004; Cavanagh *et al.* 2008; Cavanagh *et al.* 2010; Neusser et al. 2010). We did not observe a link between 6S RNA sensitivity and an extended −10 or weak −35 promoter sequence as reported in (Cavanagh *et al.* 2008) (Figure S11), nor with any other sequence features. Instead, during stationary phase, 6S RNA sensitivity was associated with low wild-type expression (Figure 4F) and low promoter occupancy by RNA polymerase (Figure 4G, based on ChIP-chip data from (Cho *et al.* 2014)). Thus, our data support a model in which sequestration of RNA polymerase by 6S RNA primarily represses promoters that bind weakly to RNA polymerase.

**The Rsd/6S RNA double knockout shows differential expression of a distinct set of genes:** We have discovered several instances of crosstalk between 6S RNA and Rsd; apart from the fact that both sequester σ^70^ in different forms, each regulates the other’s expression, and both favor the activity of σ^38^. We therefore asked whether the double knockout of Rsd and 6S RNA showed effects distinct from the single knockouts.

We observed that in every growth phase, hundreds of genes were differentially expressed in the double knockout relative to the wild-type. This far exceeded the number of differentially expressed genes in both single knockouts, demonstrating significant crosstalk between the two regulators. Figure. 5A and B highlight genes that showed differential expression in the double knockout, but less than twofold change in both single knockouts added together; there were 1780 such genes in total. These included genes encoding DNA Gyrase and Topoisomerase I, which maintain DNA supercoiling and regulate gene expression (Peter *et al.* 2004), the nucleoid-associated proteins HU, H-NS and StpA, the global transcriptional regulators ArcA, LRP and IHF, and the small RNA chaperone Hfq. This suggests that Rsd and 6S RNA act together, in a partially redundant manner, to control global gene expression and nucleoid structure.

**Figure 5.**
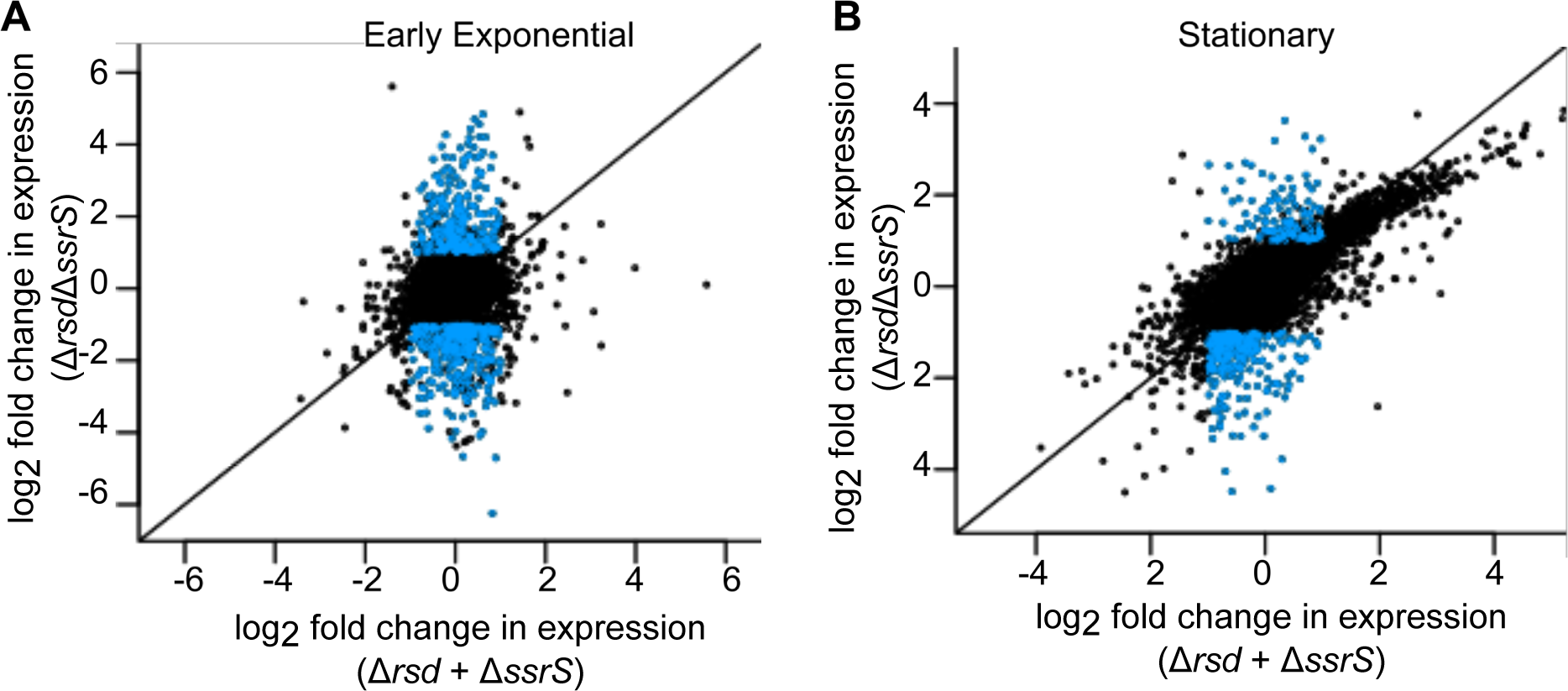
Rsd and 6S RNA have widespread non-additive effects on gene expression. Scatterplots of log2 fold change in gene expression in the Δ*rsd* Δ*ssrS* double knockout versus the sum of log2 fold changes in the Δ*rsd* and Δ*ssrS* single knockouts, for each gene, in (A) Early exponential and (B) Stationary phase. Blue points represent genes that show differential expression (>=2-fold increase or decrease, FDR-adjusted p-value<0.05) in the double knockout, but less than twofold change in expression in both single knockouts added together.

### A theoretical model suggests an explanation for the behavior of Rsd and 6S RNA

Our RNA-Seq demonstrated that Rsd and 6S RNA regulate sigma factor competition and gene expression at a global scale; however, there are two major results that appear counter-intuitive. First, how is it that Rsd, which sequesters σ^70^, increases transcription of σ^38^ target genes but has very little effect on σ^70^ target genes? Second, how does 6S RNA, which sequesters not only σ^70^, but also core RNA polymerase which is required for transcription by all sigma factors, nevertheless increase transcription by σ^38^?

To answer these questions, we constructed a mathematical model of transcription during stationary phase, using parameters from literature (Table 2). We focused on stationary phase as that is when σ^38^, 6S RNA and Rsd are at high concentrations. Our model is similar in structure to previous studies (Grigorova *et al.* 2004; Mauri and Klumpp 2014); however, it is the first to model stationary phase and to include both 6S RNA and Rsd. Consequently, our results differ from previous models.

Figure 6A shows a schematic of the model. Core RNA polymerase (E) binds to sigma factors (σ^70^ and σ^38^) forming holoenzymes (Eσ^70^ and Eσ^38^). Holoenzymes initiate transcription from target promoters (P_70_ and P_38_), releasing the sigma factor. The elongating RNA polymerase (E_e70_ and E_e38_) transcribes until released. Holoenzymes and E also bind DNA non-specifically. We focus on the steady state of this model, determined by equations (1) - (4) and (8) - (17) in Methods.

**Figure 6.**
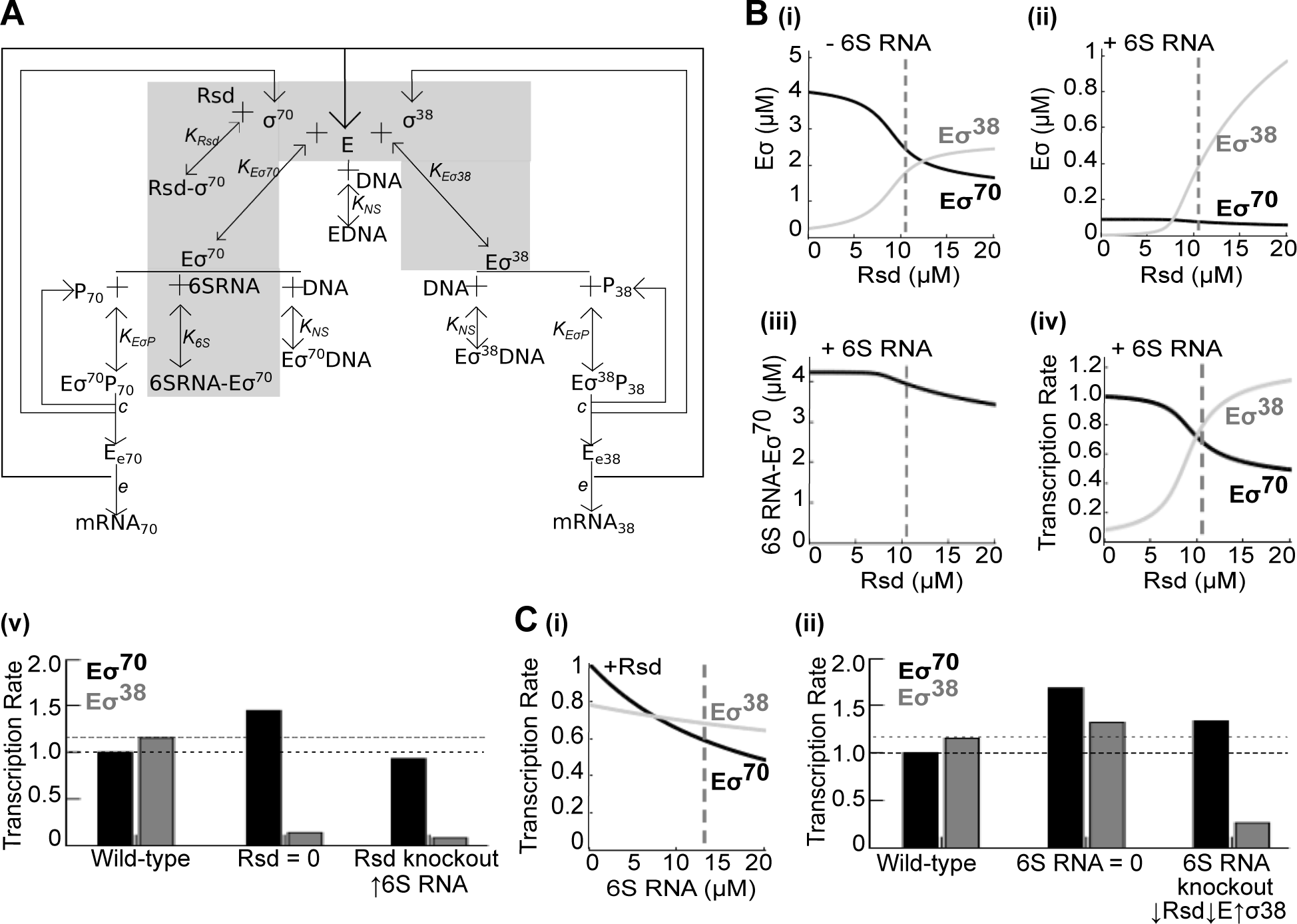
A mathematical model of Rsd and 6S RNA activity. (A) Schematic of the model. Shaded area represents reactions involved in holoenzyme formation (without DNA) (B) (i) Steady-state levels of Eσ^70^ (black) and Eσ^38^ (gray), computed from equations (1) - (9), as a function of total Rsd, when 6S RNA = 0. (ii) Same when total 6S RNA = 13 μM. (iii) Steady-state levels of the 6S RNA-Eσ^70^ complex, as a function of total Rsd. (iv) Steady-state rate of transcription from Eσ^70^ dependent promoters (black) and Eσ^38^ dependent promoters (gray), as a function of total Rsd. Vertical dashed lines represent wild-type cellular concentrations in stationary phase. (v) Steady-state rate of transcription from Eσ^70^ dependent promoters (black bars) and Eσ^38^ dependent promoters (gray bars) in the wild-type, absence of Rsd, and simulated Rsd knockout. (C) (i) Steady-state rate of transcription from Eσ^70^ dependent promoters (black) and Eσ^38^ dependent promoters (gray), as a function of total 6S RNA. Vertical dashed lines represent wild-type cellular concentrations in stationary phase. (ii) Steady-state rate of transcription from Eσ^70^ dependent promoters (black bars) and Eσ^38^ dependent promoters (gray bars) in the wild-type, absence of 6S RNA, and simulated 6S RNA knockout.

To understand how Rsd and 6S RNA regulate competition between sigma factors, we initially modeled the formation of Eσ^70^ and Eσ^38^ in the absence of DNA. This is represented by the shaded area in Figure 6A and steady-state equations (1) - (9) in Methods. Figure 6B(i) depicts what happens when Rsd is added to a system containing only E, σ^70^ and σ^38^. Each value of Rsd on the x-axis corresponds to a separate ’run’ of the model where we compute the steady-state for those parameter values, before moving on to the next Rsd value. We predict that as Rsd is increased, it sequesters σ^70^, reducing Eσ^70^ formation. This allows more E to bind σ^38^, increasing the formation of Eσ^38^, consistent with previous predictions of Rsd function (Jishage and Ishihama 1999).

However, Figure 6B(ii) predicts that when 6S RNA is also present, Rsd increases Eσ^38^ with little effect on Eσ^70^. How? This paradoxical result can be understood with Figure 6B(iii), which shows that the concentration of the 6S RNA-Eσ^70^ complex decreases as Rsd is increased. When 6S RNA is present, increasing Rsd still reduces E-σ^70^ association; however, the reduction in Eσ^70^ is compensated for by the release of Eσ^70^ from its complex with 6S RNA, and so there is little change in the overall Eσ^70^ level. This prediction remains when we include DNA in the model (represented by the complete schematic in Figure 6A). Figure 6B(iv) shows that simulating increased Rsd increases transcription by Eσ^38^ with less effect on Eσ^70^ transcription.

Our Δ*rsd* strain also displayed a ~2.3-fold increase in 6S RNA during stationary phase. Figure 6B(v) shows the predicted rate of transcription from Eσ^70^ and Eσ^38^ target promoters in the wild-type and when Rsd = 0. The third pair of bars is an approximation of conditions in the Δ*rsd* strain, where 6S RNA is increased 2.3-fold. We see that increased expression of 6S RNA could reduce Eσ^70^ dependent transcription in the Δ*rsd* strain to almost the wild-type level, so that the main observable effect of knocking out Rsd would be reduced transcription by Eσ^38^.

This explains why, in both our RNA-seq and a previous study^27^, Rsd increases transcription of Eσ^38^ target genes, with little to no effect on Eσ^70^. We hypothesize that high levels of 6S RNA cause Rsd to act as a σ^38^ regulator and not a σ^70^ regulator, and, moreover, that Rsd controls the level of 6S RNA to modulate its own function.

We also observed a 2.4-fold increase in *oxyS* (a repressor of *rpoS* translation) in the Rsd knockout. We did not include this in the model due to lack of quantitative data on *oxyS;* however, it is possible that a small reduction in σ^38^ protein levels, perhaps due to increased *oxyS*, may contribute to reduced σ^38^ activity in the Δ*rsd* strain. However, it is difficult to confirm this.

What is the effect of 6S RNA? Our model predicts that as 6S RNA levels increase, it sequesters Eσ^70^, reducing the amount of E available to bind to both sigma factors and thus inhibiting formation of both holoenzymes. This decreases the rate of transcription by both holoenzymes (Figure 6C(i)). However, our RNA-seq showed that deleting 6S RNA actually results in reduced transcription of Eσ^38^ target genes, i.e. 6S RNA increases transcription by Eσ^38^. This is also supported by single-promoter studies in (Trotochaud and Wassarman 2004). How is this possible? A 10-fold increase or decrease in any of the parameters was not sufficient to reproduce this observation.

Apart from losing 6S RNA, our Δ*ssrS* strain also displayed reduced Rsd and RpoB, and increased σ^38^. Figure 6C(ii) shows the predicted rate of transcription from Eσ^70^ and Eσ^38^ target promoters in the wild-type and when 6S RNA = 0. The third pair of bars is an approximation of conditions in the Δ*ssrS* strain; Rsd and E are reduced by 50% and σ^38^ is increased by 50%. Eσ^38^ transcription is now reduced to less than the wild-type level; in fact, within the default parameters of our model, reducing Rsd alone from 10.4 μM to 8 μM is sufficient to lower Eσ^38^ transcription in the 6S RNA knockout below its wild-type level.

We also observed increased *crl* mRNA in Δ*ssrS*. We have not modeled this due to lack of quantitative data on Crl. However, if the change in mRNA corresponds to increased Crl protein, it could partially mitigate the effect of reduced Rsd in the Δ*ssrS* strain.

Thus, from our model we can make the strong claim that the binding reactions of Rsd and 6S RNA shown in Figure 6A cannot explain the reduced transcription of σ^38^ target genes in the Δ*ssrS* strain. However, adding reduced Rsd levels to the model is sufficient. We therefore hypothesize that the reduced Eσ^38^ transcription in the 6S RNA knockout is due to its indirect effects primarily via Rsd.

The quantitative predictions of this model would ideally be validated in detail by *in vitro* transcription experiments, measuring the change in transcription of σ^38^ and σ^70^ dependent promoters with Rsd and 6S RNA in varying concentrations. We predict that the effect of Rsd on σ^70^ dependent transcription should decrease as 6S RNA is increased. We also predict that increasing 6S RNA alone should reduce both σ^38^ and σ^70^ dependent transcription, but σ^38^ dependent transcription can be increased on increasing Rsd. To our knowledge, such studies have not been carried out. However, such experiments are necessarily limited to a few promoters. Some predictions may also be testable by looking at distributions of gene expression *in vivo*; for instance, increasing Rsd expression in Δ*ssrS* back to the wild-type level should largely mitigate the reduced expression of σ^38^ target genes.

### DISCUSSION

Rsd and 6S RNA have long been known to regulate RNA polymerase in *E. coli.* Here we report for the first time that Rsd regulates gene expression from early exponential to stationary phase. Though it sequesters σ^70^, its primary function is reducing transcription by the alternative sigma factor σ^38^. Based on theoretical modeling, we suggest that this is due to 6S RNA, which minimizes the effect of Rsd on Eσ^70^ levels. We show that Rsd regulates 6S RNA expression, thereby minimizing its own effect on Eσ^70^. Since Rsd overexpression increases transcription by σ^24^ and σ^54^ in a ppGpp^0^ background (Laurie *et al.* 2003; Costanzo *et al.* 2008), Rsd may generally assist alternative sigma factors to associate with RNA polymerase, under suitable conditions.

6S RNA is more complex, regulating expression of >400 genes; however, we did not observe associations between promoter sequence and 6S RNA susceptibility as previously reported (Cavanagh *et al.* 2008). Our data is more similar to that of (Neusser *et al.* 2010), who reported reduced expression of *rpoB* and ribosomal genes in Δ*ssrS* in stationary phase, and increased ppGpp without increase in *relA* (*spoT* levels were found unchanged in all studies). However, there is still relatively low overlap in the list of 6S RNA regulated genes (46 genes). This is likely because of differences in time points and media. Even within our dataset, there is little overlap between genes regulated by 6S RNA at different time points. It seems that the 6S RNA regulon varies greatly under different conditions. This supports a model where 6S RNA acts as a background-level regulator operating on RNA polymerase, with gene-level outcomes depending strongly upon cellular environment; this could involve DNA topology, transcription factors, and other RNA polymerase-binding factors, all of which vary with growth phase (Dorman 2013). Regulators that interact with 6S RNA can be experimentally identified, for example by transposon insertion sequencing to identify genes whose disruption leads to a larger survival defect in a 6S RNA knockout than in a wild-type background, or by *in vitro* transcription experiments testing whether the ability of 6S RNA to regulate known targets is affected by the presence of other regulators in the medium.

Previous work using individual promoters (Trotochaud and Wassarman 2004) had suggested that 6S RNA might increase transcription by Eσ^38^ during stationary phase. It was unknown whether this was a direct effect of 6S RNA, allowing σ^38^ to compete more effectively for E, or indirect, via a trans-acting factor important for σ^38^ activity. However, another study (Neusser *et al.* 2010) found no evidence of this link. We show that 6S RNA increases transcription by Eσ^38^ globally, from early exponential to stationary phase, and show mathematically that this is extremely unlikely to be a direct effect of sequestration. We show that 6S RNA increases Rsd protein level and suggest that Rsd is primarily responsible for 6S RNA’s effect on Eσ^38^ activity.

It has been asked, given 6S RNA’s function as a global regulator, why its deletion does not cause a growth defect. Our data, along with others (Cavanagh *et al.* 2010; Neusser *et al.* 2010), shows multiple feedback effects in the Δ*ssrS* strain, where the cell reduces RNA polymerase expression to compensate for 6S RNA loss, and increases σ^38^ and possibly Crl to compensate for reduced Eσ^38^ activity. Apart from these two, increased ppGpp may also have such a compensatory role during TS phase, while in LS phase, the 6S RNA knockout has higher expression of *ppk*, responsible for synthesis of polyphosphate - which activates transcription by Eσ^38^ (Kusano and Ishihama 1997). In stationary phase, the Rsd/6S RNA double knockout shows reduced expression of DNA supercoiling enzymes; supercoiling regulates promoter binding by Eσ^70^ and Eσ^38^ (Kusano *et al.* 1996; Bordes *et al.* 2003); in ME phase, it shows reduced expression of Hfq, which increases σ^38^ expression (Zhang *et al.* 1998). These relationships, illustrated in Figure S12, demonstrate that competition between sigma factors is very tightly controlled, and 6S RNA is connected to multiple pathways involved in this process. We see that 6S RNA and Rsd regulate each other’s expression, and control each other’s effect on gene expression; finally, we report that 1780 genes across 5 growth phases - almost 40% of the genes in the cell - are differentially expressed in the 6S RNA-Rsd double knockout but not in the single knockouts added together. This provides direct evidence that 6S RNA and Rsd have non-additive effects on gene expression, since the single knockouts cannot explain the widespread gene expression changes observed in the double knockout. Previous studies have only examined single knockouts; this underscores the importance of studying 6S RNA and Rsd as a unit that controls global RNA polymerase activity.

Given that 6S RNA homologs are widespread in bacteria, co-occurring with Rsd and with other RNA polymerase regulators such as the actinobacterial RbpA, and many species have two or three 6S RNA homologs with different expression patterns and potentially different binding partners (Barrick *et al.* 2005; Wehner *et al.* 2014), we suggest that studying the relationships between RNA polymerase regulators will give greater insights into transcriptional control across the bacterial kingdom.

## ACKNOWLEDGMENTS

Illumina sequencing was performed at the Centre for Cellular and Molecular Platforms. Strains and plasmids were obtained from CGSC. We thank Revathy Krishnamurthy and Harshvardhan for assistance with experiments. We also thank Prof. Dipankar Chatterji, Prof. Akira Ishihama, Prof. Karen Wassarman, Prof. Steve Busby, Dr. Dasaradhi Palakodeti, and Prof. M.M. Panicker for discussions.

